# Sex-based mutational landscape confers sensitivity to checkpoint blockade immunotherapy in cutaneous melanoma

**DOI:** 10.1101/868950

**Authors:** Mingming Jia, Tian Chi

## Abstract

Anti-programmed cell death protein 1 (PD-1) therapy provided superior benefits in skin cutaneous melanoma (SKCM), but only a minority of patients responded. As SKCM had sex-based immunological differences, we sought to explore the potential interactions between sex and clinical responses to anti-PD-1 therapy in SKCM. We found that sex had significant effects on anti-PD-1 therapy. Tumor mutation burden (TMB) and neoantigen burden significantly correlated with the clinical responses to anti-PD-1 therapy only in males. Meanwhile, we recruited The Cancer Genome Atlas (TCGA) database to explore sex-based differences of the tumor microenvironment (TME) in SKCM. We observed that males with high TMB, especially in conjunction with interferon-gamma (IFN-γ) signaling, significantly correlated with increased PD-L1 expression, major histocompatibility complex (MHC) class I gene expression, and the infiltration of cytotoxic T lymphocytes (CTLs). In addition, TMB and anti-PD-1 efficacy in SKCM each correlated with homologous recombination repair (HRR) gene BRCA2 mutations, but only in males. Taken together, these data revealed that in SKCM, high TMB correlated with prominent immunotherapeutic TME only in males, and that gender should be taken into account when predicting the efficacy of anti-PD-1 therapy for SKCM.

## Introduction

Sex-based immunological differences have been reported across multiple cancer types, with males getting higher risks of mortality from malignant cancers than females (1, 2). Notably, melanoma displays one of the greatest sex-specific outcomes among many cancers (2, 3). These sex-based differences of tumorigenesis in melanoma probably reflect genetic, hormonal, environmental, and microbial effects on the immune system throughout life in humans (1, 4–6).

Recent clinical trials have demonstrated promising performance on PD-1 immune checkpoint blockade (ICB) therapies in melanoma, but only a fraction of patients responded (7–11). Previous reports indicated that tumor mutation burden (TMB) and neoantigen burden (12, 13), PD-L1 expression (12, 14), interferon-gamma (IFN-γ)-related signaling (7, 15, 16), major histocompatibility complex (MHC) gene expression (10, 17), the abundance of cytotoxic T lymphocytes (CTLs) infiltration (18, 19), and mismatch repair (MMR) deficiency (14) could be predictive biomarkers for ICB therapies across multiple cancer types (14, 20). Among these biomarkers, TMB performs much better and is being developed as an immunotherapeutic biomarker for the oncology clinic (14, 21, 22). High TMB could potentially produce more immunogenic neoantigens on the surface of tumor cells, and T cells were then activated to proliferate and destroy these tumor cells presenting the recognized neoantigens (14, 21, 22). However, the predictive power of TMB for ICB response is still imperfect and needs further improvement (14, 21, 22).

Previous studies have reported that increased TMB was associated with clinical efficacy to cytotoxic T lymphocyte antigen 4 (CTLA-4) blockade but not PD-1 blockade in melanoma (8, 13, 18), and male patients had higher TMB than females in melanoma (23, 24). Consequently, we hypothesized that male factor integrating with known biomarkers, especially TMB, could provide more accurate predictive power of ICB response in skin cutaneous melanoma (SKCM), one of the commonest subtypes of melanoma (25). Although a recent meta-analysis concluded that males could derive greater benefits from ICB therapies than females in many cancer types (23), the conclusion was controversial (26–28). In addition, their work also did not consider and analyze the sex-based association of TMB with the efficacy of immunotherapies.

To test our hypothesis, we captured two well-studied melanoma immunotherapeutic cohorts (8, 9) and one non-immunotherapeutic The Cancer Genome Atlas (TCGA) dataset. It has already been indicated that many public tumor databases without immunotherapies were still informative regarding tumor immune evasion (11, 19, 29). Therefore, we fully integrated these three cohorts to comprehensively analyze interactions between sex-based biomarkers and the efficacy of immunotherapies. Significantly, we uncovered that males with high TMB, in conjunction with other biomarkers, could be a superiority group to anti-PD-1 therapy in SKCM.

## Results

### Clinical and genomic characteristics of the study cohorts

After extensive survey of cancer databases and literature, we were able to collect only two melanoma PD-1 blockade cohorts with both clinical and genomic characteristics: 34 SKCM patients in Hugo et al study (8) and 44 SKCM samples in Riaz et al study (9). Pre-therapy biopsies from SKCM patients were determined by whole-exome sequencing (WES). Given the small number of samples in these cohorts, we integrated the data into one Cell-SKCM dataset according to their common features, such as patients’ TMB, neoantigens, gender, clinical response to anti-PD-1 therapy, histology, and age (n=78; Fig. 1A and B; Table S1). In addition, we also recruited TCGA-SKCM datasets containing multidimensional non-immunotherapeutic characteristics to further explore the tumor immune escape in Cell-SKCM cohort (n=472; Fig. 1C; Table S1). Importantly, in Cell-SKCM, the quantity and range of TMB (i.e., the total number of somatic non-synonymous single nucleotide variations) were similar to that in the TCGA-SKCM dataset (*P*=0.71; Fig. S1A), suggesting the homogeneity of these two cohorts as previously described (12, 30).

**Figure 1.**
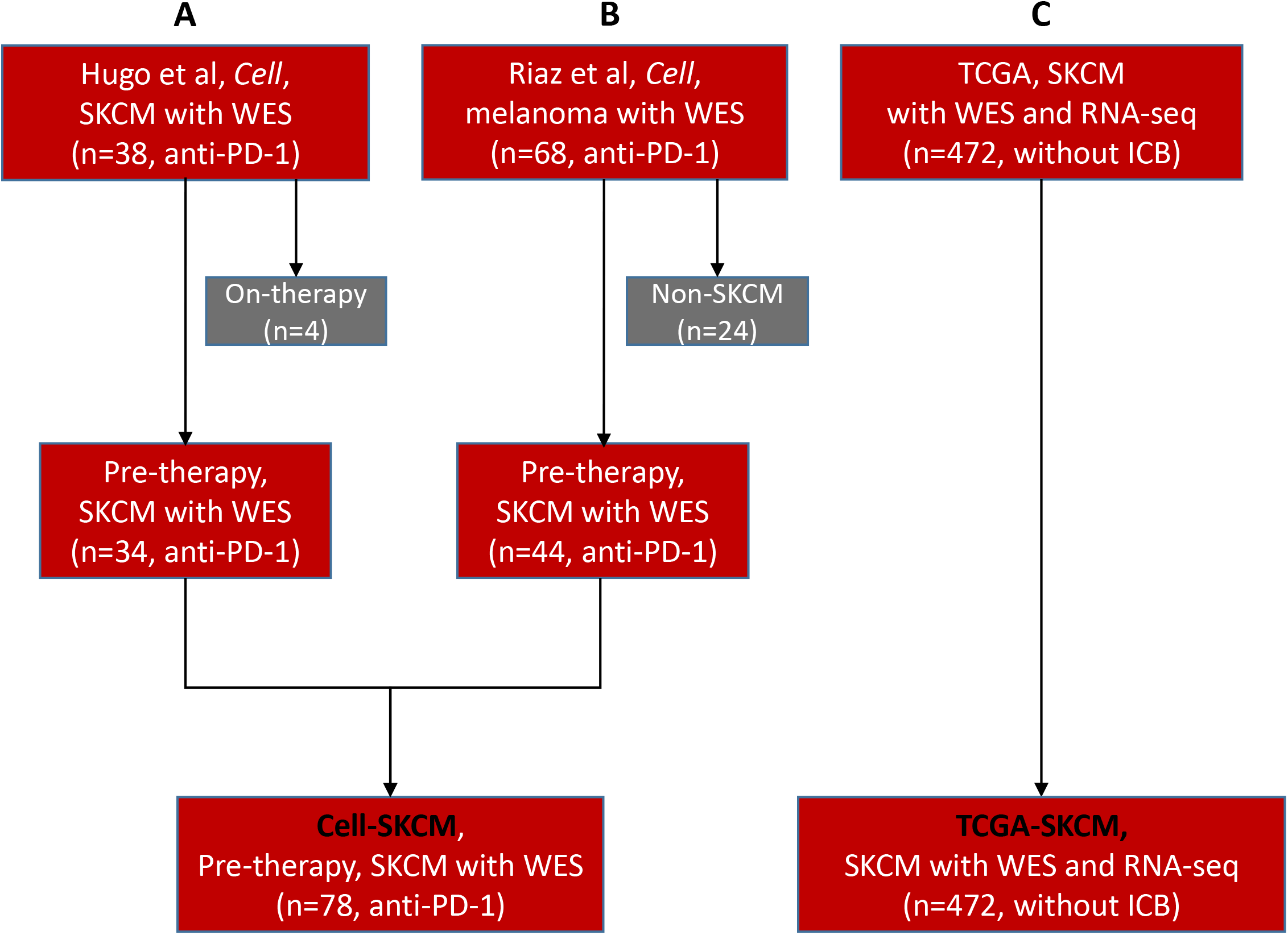
Study design. (A and B) two PD-1 blockade immunotherapeutic cohorts published on *Cell* were integrated into a single Cell-SKCM cohort. Pre-therapy biopsies from patients were assessed by WES before treatment with the PD-1 blockade therapy. We abandoned those samples that were on-therapy or non-SKCM. (C) One TCGA non-immunotherapeutic SKCM dataset with both WES and RNA-seq data available was defined as the TCGA-SKCM cohort. SKCM, skin cutaneous melanoma; WES, whole-exome sequencing; TCGA: the cancer genome atlas. ICB, immune checkpoint blockade.

### Sex differences correlated with TMB and neoantigen burden in patients with differential response to anti-PD-1 therapy

Compared with females, TMB was markedly increased in males in the TCGA-SKCM cohort; a similar trend was observed for the Cell-SKCM cohort although the difference was not statistically significant possibly because of smaller sample size (Fig. S1B). Consistent with these, males but not females displaying good therapeutic responses (CR/PR: complete/partial response to anti-PD-1 therapy) showed significantly higher TMB compared with those who had no response (SD/PD: stable/progressive disease for anti-PD-1 therapy) in Cell-SKCM cohort (Fig. 2A). Receiver operator characteristic (ROC) curves also validated high associations between TMB and clinical efficacy to PD-1 blockade in males (AUC=0.74, *P*=0.005; Fig. 2B). The discovery of the sex-based immunotherapeutic differences of TMB was consistent with neoantigen burden (AUC=0.73, *P*=0.007; Fig. 2D and E); neoantigens are generated as a consequence of TMB and can elicit effective immune responses (12, 31).

**Figure 2.**
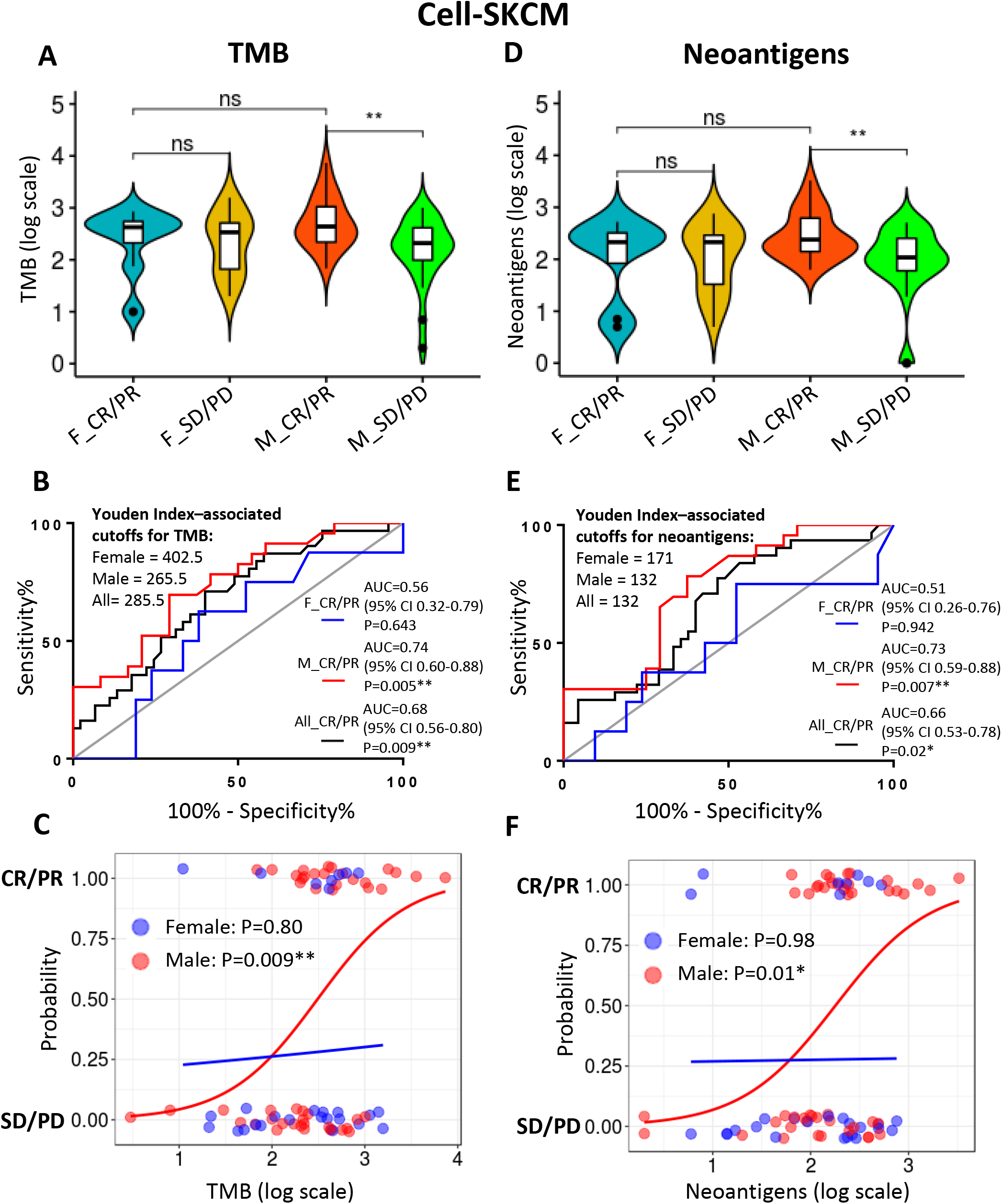
Correlations of sex differences with TMB and neoantigens in patients with differential responses to anti-PD-1 therapy. (A and D) Sex-based differences of TMB and neoantigens in Cell-SKCM cohort with differential responses to anti-PD-1 therapy. (B and E) Sex-based differences of ROC curves for correlations of TMB and neoantigens with CR/PR to anti-PD-1 therapy. Youden Index–associated cutoffs for TMB and neoantigens in each group are shown. The area under the ROC curve (AUC); CI (confidence interval). (C and F) The logistic regression of TMB and neoantigens with clinical response to anti-PD-1 therapy. The S-shaped curves in males are clearly better fits for most of the data than in females. Probability, probability of getting clinical benefits from anti-PD-1 therapy. F_CR/PR (n=8), females with complete/partial response; F_SD/PD (n=21), females with stable/progressive disease; M_CR/PR (n=23), males with complete/partial response; M_SD/PD (n=24), males with stable/progressive disease; All_CR/PR (n=31), all samples with complete/partial response.

To further reveal the potential clinical usefulness of TMB and neoantigen burden for predicting PD-1 blockade response, we used the Youden index (7, 11, 12) to determine the optimum cutoff point. The cutoff points of both TMB and neoantigens were significantly lower in males than in females, suggesting that these two biomarkers were more sensitive to males for predicting PD-1 blockade efficacy (Fig. 2B and E). Additionally, univariate logistic regression analysis also demonstrated that TMB and neoantigens were significantly associated with clinical efficacy to anti-PD-1 therapy in male patients (Odds Ratio=7.96, *P*=0.009 and Odds Ratio=8.01, *P*=0.01, respectively) (Fig. 2C and F; Table S2). Using multivariate logistic regression analysis with adjustment for age, we also observed a significant association between male-based TMB and clinical efficacy to anti-PD-1 therapy (Odds Ratio=7.77, *P*=0.01; Table S2). Of note, given its significant correlation with TMB, neoantigens were not included in the multivariate regression analysis (Fig. S2A and B).

To assess the clinical utility of TMB in Cell-SKCM cohort, patients were classified into four TMB-defined groups based on Youden index-associated cutoff points (11): females with high TMB (F_High), ≥ 402.5; females with low TMB (F_Low), < 402.5; males with high TMB (M_High), ≥ 265.5; males with low TMB (M_Low), < 265.5. The same methodology applied to neoantigen-defined groups. This stratification scheme revealed that the percentage of overall survival (OS) was significantly better in males with high TMB and neoantigen burden (*P*=0.03 and *P*=0.05, respectively; Fig. S3A-D).

### Correlations of sex differences with interferon-γ (IFN-γ) signaling and related gene signatures in SKCM

As Cell-SKCM cohort lacks transcriptomic profile, we turned to multidimensional TCGA-SKCM datasets, which carry both genomic and matched transcriptomic profiles, to further explore the sex-based tumor immune evasion. We stratified the TCGA-SKCM samples into four TMB-defined groups as described above for the Cell-SKCM cohort (Fig. S1C; Table S3), observing significant sex-based differences in neoantigens in different groups, consistent with a tight correlation between TMB and neoantigens (Fig. S2C and D).

Furthermore, males (but not females) with high TMB were significantly enriched in genes associated with immune system and IFN-γ signaling (such as PD-L1, IFN-γ, GBP1, HLA-B, HLA-C, and CTLA4; Fig. 3A-C), a finding reinforced by unsupervised hierarchical clustering of differential expression genes revealing that only males with high TMB significantly increased expression of genes related to IFN-γ signatures (such as IFN-γ, STAT1, CXCL9, CXCL10, CXCL11, and IDO1; Fig. 4A-C). Our finding supports the notion that IFN-γ signaling and related gene signatures play important roles in mediating tumor rejection for ICB therapies and can predict clinical response to PD-1 blockade in melanoma (7, 10, 15, 16). Finally, a T cell-inflamed gene expression profile (GEP) has been described for predicting PD-1 blockade response across multiple cancers (7, 11). Indeed, T cell-inflamed GEP score tended to increase in males with high TMB (*P*=0.07; Fig. S4A). The low statistical significance might be attributed to the fact that T cell-inflamed GEP (as a pan-tumor biomarker) is not as specific and effective as IFN-γ signatures in predicting PD-1 blockade efficacy in SKCM (7).

**Figure 3.**
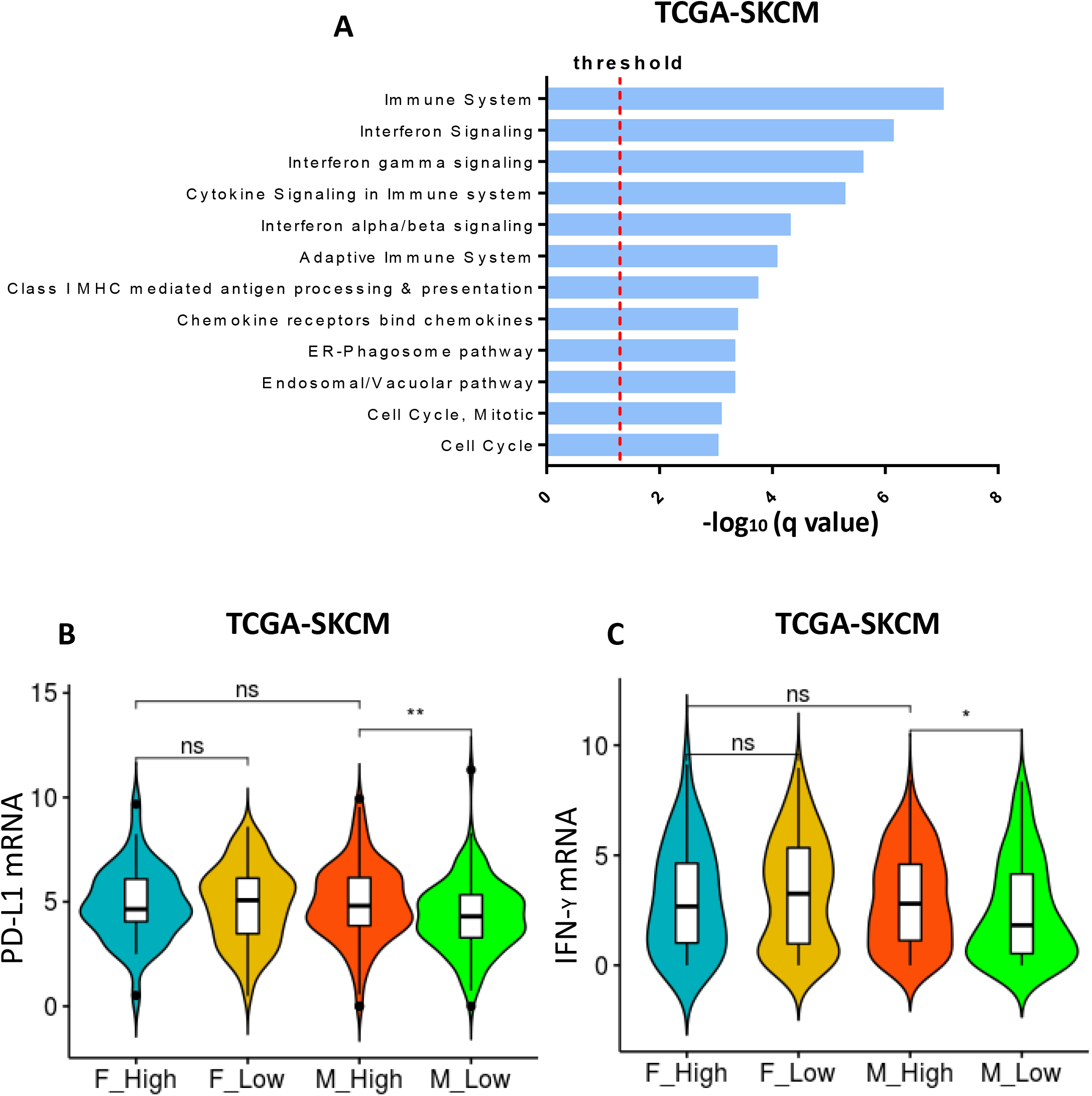
Correlations of sex differences with interferon-γ (IFN-γ) signaling in TCGA-SKCM dataset with differential TMB. (A) Pathway enrichment analysis of differential expression genes (log2 fold change ≤0.5 and *P* value ≤0.05) between males with high TMB and low TMB (pathways with q value ≤0.05 significant). (B and C) Quantitative analysis of mRNA expression of PD-L1 and IFN-γ. F_High (n=39); F_Low (n=141); M_High (n=169); M_Low (n=123).

**Figure 4.**
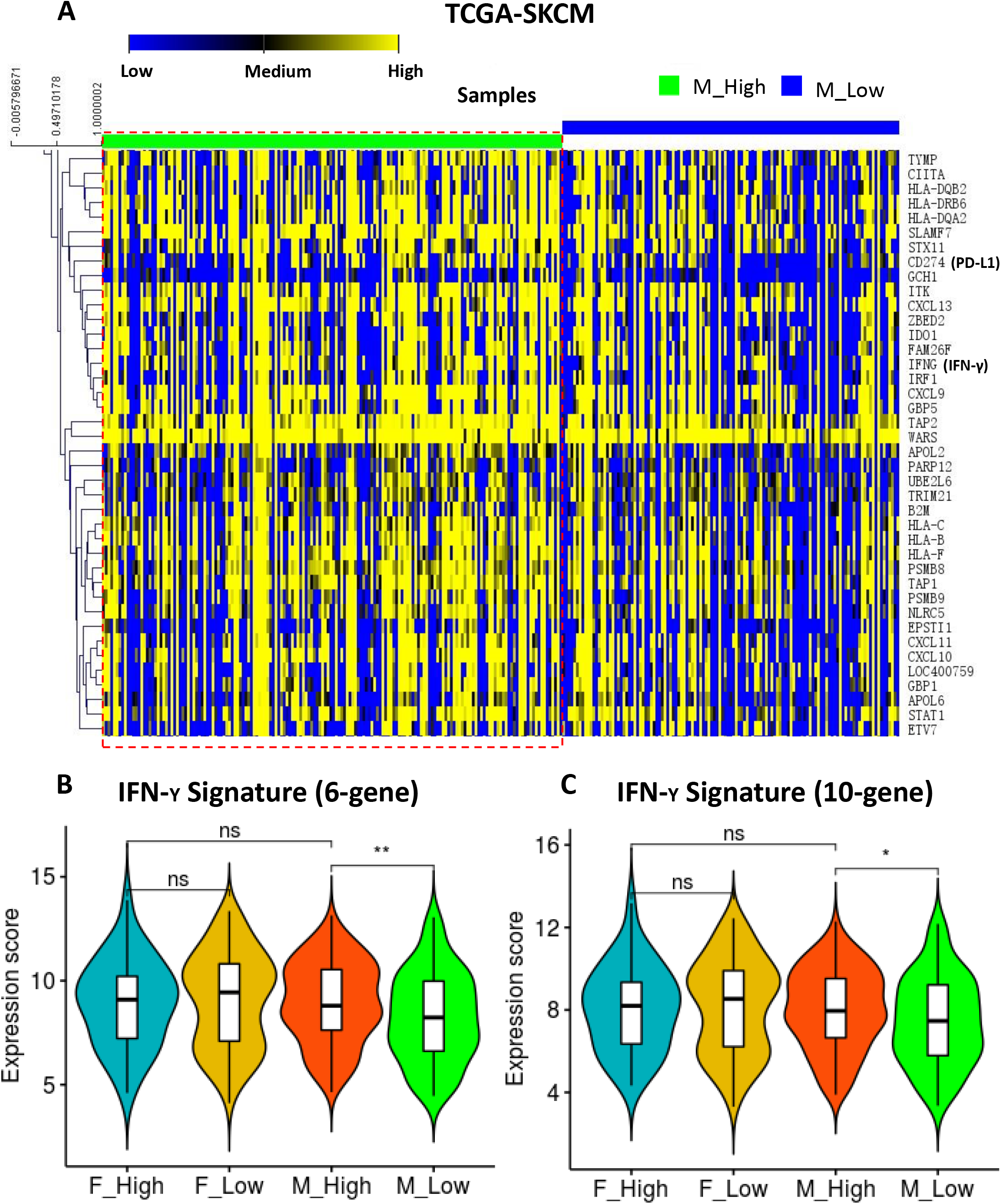
Correlations of sex differences with IFN-γ-related gene signatures in TCGA-SKCM with differential TMB. (A) Hierarchical clustering of differentially expressed genes between males with high TMB and low TMB (differentially expressed genes with *P* value ≤0.05). (B and C) Quantitative analysis of two IFN-γ signatures associated with clinical response to anti-PD-1 therapy among four groups. F_High (n=39); F_Low (n=141); M_High (n=169); M_Low (n=123).

### IFN-γ-induced immune microenvironment facilitates MHC-I gene expression in males with high TMB

IFN-γ-mediated signaling can upregulate or induce the expression of MHC-I molecules on non-immune cells in tumor tissues, which can accelerate immune recognition and removal of malignant cells (16). Additionally, the reduction in the MHC-I gene copy number can facilitate immune evasion via defects in neoantigen presentation (32), and patient MHC-I molecules can influence cancer response to ICB therapies (10, 17). Consequently, we next examined MHC-I gene expression in the TCGA-SKCM dataset. Unsupervised clustering of differentially expressed genes in TCGA-SKCM samples showed that MHC-I genes (such as HLA-A, HLA-B, and HLA-C) were significantly enriched in males but not females with high TMB (Fig. 4A), concordant with the pathway analysis (Fig. 3A). MHC-I gene copy numbers were similar among the four groups (Fig. 5A, C, and E), but MHC-I gene expression was highly upregulated in males with high TMB (Fig. 5B, D, and F), presumably as a result of IFN-γ stimulation which should facilitate antigen presentation and T cell recognition in the tumor microenvironment.

**Figure 5.**
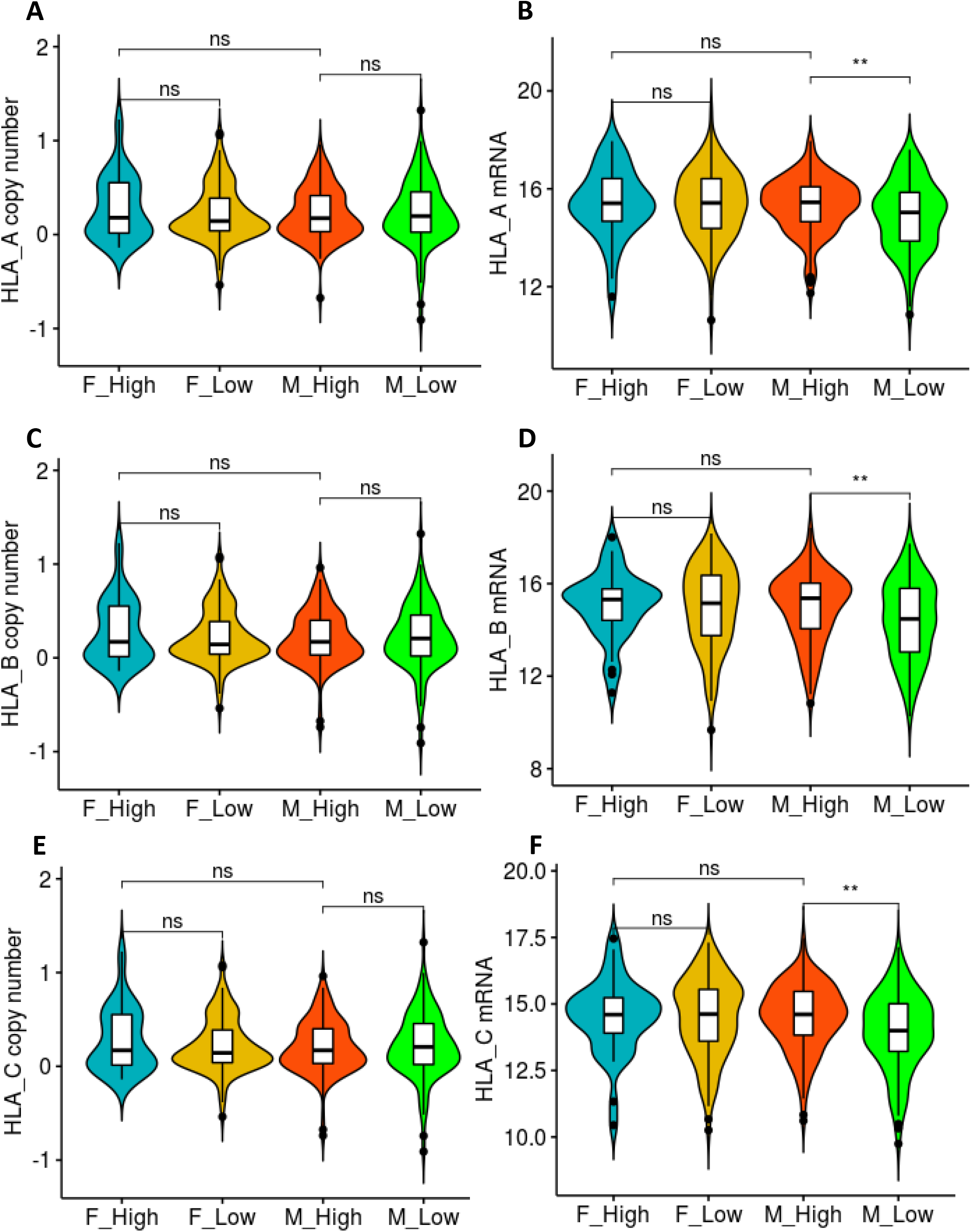
Correlations of sex differences with MHC-I gene expression in TCGA-SKCM with differential TMB. (A, C, and E) Quantitative analysis of MHC class I gene copy number variations (CNVs) among four groups. (B, D, and F) Quantitative analysis of MHC class I gene mRNA expression among four groups. F_High (n=39); F_Low (n=141); M_High (n=169); M_Low (n=123).

### CTLs infiltration in IFN-γ-induced immune microenvironment

The presence of cytotoxic T lymphocytes (CTLs) is an important biomarker for predicting response to ICB therapies (18, 19). We therefore examined the crucial markers of CTLs (GZMA, GZMB, CD8A, CD8B, and PRF1) as described (19), finding that CTLs infiltration tended to increase in males with high TMB compared with low TMB (*P*=0.21; Fig. S4B).

CTLs infiltration can be affected by its dysfunctional state and immunosuppressive factors (19). Using Tumor Immune Dysfunction and Exclusion (TIDE) algorithm (19), we found that males with high TMB displayed significantly decreased exclusions of T cells (*P*=0.0067) but no T cell dysfunction, suggesting that the infiltrating CTLs were functional and thus beneficial to these patients (Fig. 6A and B). Consistent with this, these patients also showed decreased TIDE scores and increased IFN-γ scores (*P*=0.0069 and *P*=0.0059, respectively; Fig. 6C and D), which are known to correlate with the efficacy of anti-PD-1 therapy.

**Figure 6.**
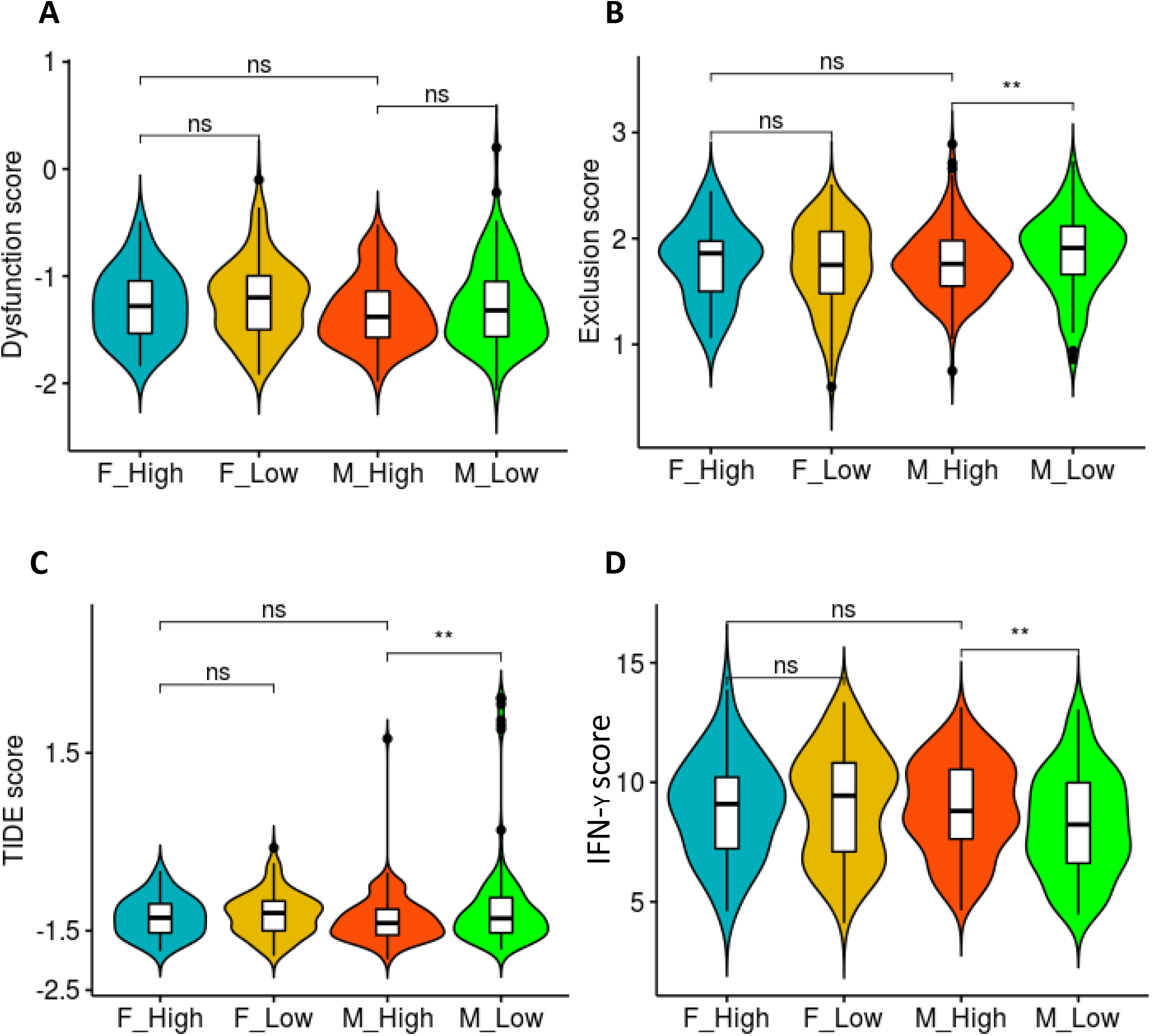
Correlations of sex differences with the infiltration of CTLs in TCGA-SKCM with differential TMB. (A and B) Relative quantification of T cell dysfunction and exclusion among four groups. (C and D) Relative quantification of TIDE and IFN-γ score among four groups. F_High (n=39); F_Low (n=141); M_High (n=169); M_Low (n=123).

### BRCA2 mutation correlated with therapeutic response only in males

Melanoma is one of the most typical mutation-caused malignant tumors and shows a higher prevalence of C>T mutations through ultraviolet exposure and other lesions (33). We found that both males and females with high TMB showed a higher proportion of C>T mutations, suggesting no gender bias in ultraviolet-induced mutation signature (Fig. S5A and B).

In contrast, Gene Set Enrichment Analysis (GSEA) showed prominent enrichment of pathways associated with cell cycle and DNA replication only in males with high TMB (Fig. 3A; Fig. 7A and B), which could potentially facilitate the accumulation of DNA mutation and damage.

**Figure 7.**
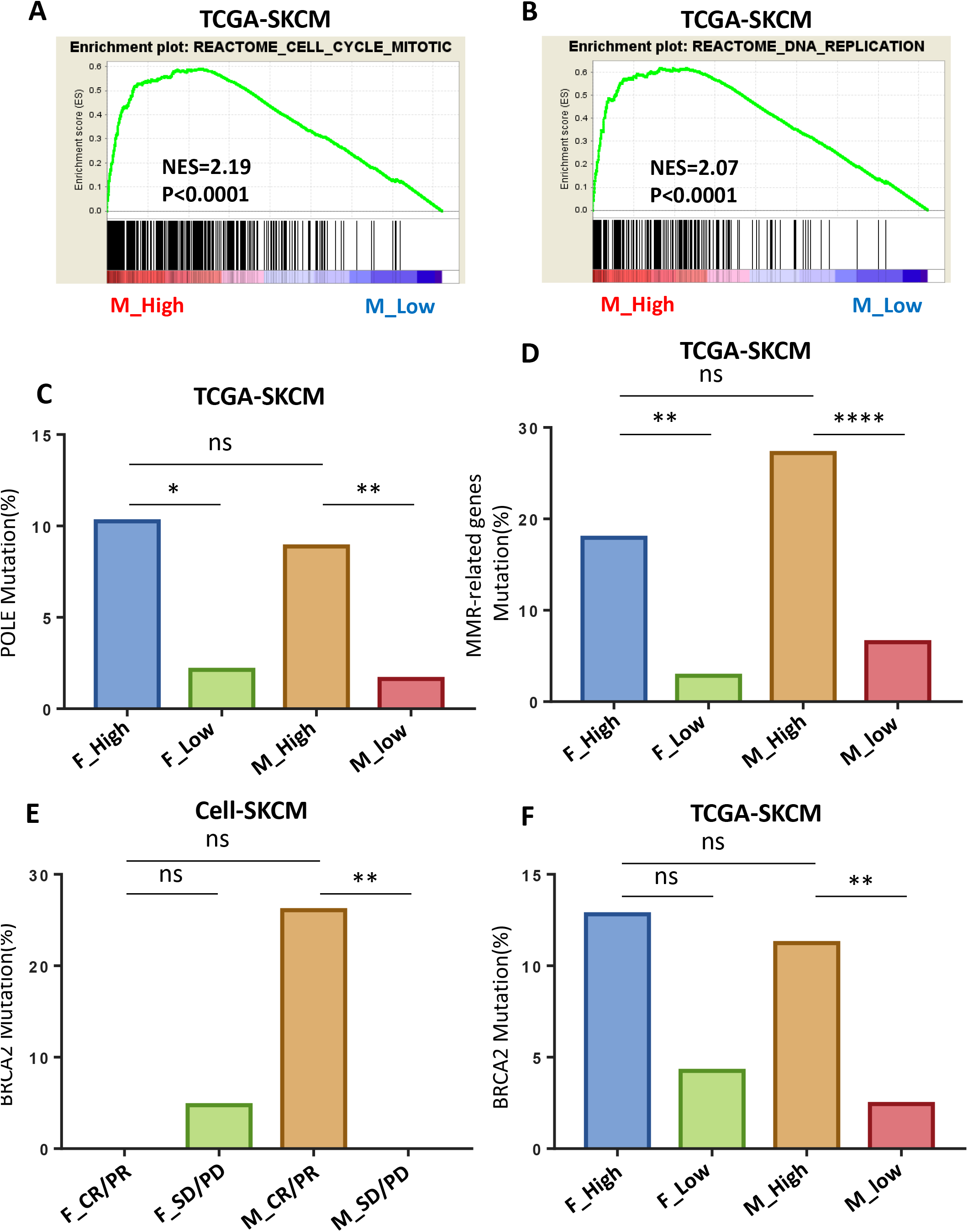
Correlations of sex differences with DDR pathways in TCGA-SKCM with differential TMB. (A and B) Males with high TMB upregulated genes involved in cell cycle and DNA replication pathways. NES, normalized enrichment score. (C and D) Mutation percentage of POLE and MMR-related genes among the four groups in TCGA-SKCM. (E) Males who responded to anti-PD-1 therapy showed significantly increased mutations of BRCA2 in Cell-SKCM cohort. (F) Males with high TMB showed significantly increased mutations of BRCA2 in TCGA-SKCM cohort. F_High (n=39); F_Low (n=141); M_High (n=169); M_Low (n=123).

We further examined the DNA damage response (DDR) pathways essential for genomic integrity (34), focusing on the genes whose mutations can induce a hypermutational phenotype and durable clinical benefits from ICB therapies, such as base excision repair (POLE), MMR (MSH2, MSH6, ARID1A, MLH1, and PMS2) and HRR (BRCA2) (34, 35). We found that mutations in POLE and MMR-related genes were increased in both females and males with high TMB (Fig. 7C and D). However, BRCA2 mutations were specifically enriched in males with clinical benefits from anti-PD-1 therapy (*P*=0.009) and with high TMB (*P*=0.006), indicating that males who had high TMB and BRCA2 mutations were more sensitive to anti-PD-1 therapy in SKCM (Fig. 7E and F). Furthermore, to assess the biologic and oncogenic effects of these genetic alterations, we performed the analysis with MutationMapper (36), PolyPhen-2 (37), and SIFT (38). The result revealed that most of these genetic mutations (POLE, MMR-related genes and BRCA2) were loss-of-function in nature (results of MutationMapper, PolyPhen-2, and SIFT not shown).

### Correlations of estrogen and estrogen receptors (ERs) with tumor immune escape in SKCM

We have found that only in males was high TMB correlated with other known biomarkers for predicting PD-1 blockade efficacy in SKCM (Fig. S6A and B). To begin to explore the molecular basis of this gender-specific phenomenon, we focused on ERβ, the predominant subtype of ERs in melanoma (39). ERβ acts as a tumor suppressor (39) partly via promoting IFN-γ secretion in the TME (1, 40), and its expression decreases as melanoma progresses. We noted a marked decrease in ERβ expression in females with high TMB compared with males with high TMB (Fig. S7A and B), and there were tighter associations between ERβ and IFN-γ expression in females with high TMB than males with high TMB (r=0.24, r= −0.03, respectively; Fig. S7C). Thus, in females with high TMB, ERβ and IFN-γ expression were compromised, which might help disrupt the correlation of TMB and other biomarkers with the therapeutic efficacy, and this disruption may be exacerbated after menopause, when the estrogen levels decline rapidly (41). Of note, in the female SKCM patients, PD-L1 expression and IFN-γ score also decreased with age, which could further disrupt the above-mentioned correlation (Fig. S8A-D).

## Discussion

Very little is known about sex-associated differences for ICB therapies. Although a recent meta-analysis reported that males could derive greater value from ICB therapies than females in multiple cancers (23), their conclusion is controversial (26–28). For instance, Wallis and colleagues’ meta-analysis collecting more clinical trials showed no statistically significant differences between patient sex and immunotherapy efficacy in many advanced cancers (26). Another recent study also reported no significant association of sex with the efficacy of immunotherapies in metastatic renal cell carcinoma (28). A caveat of these studies is that anti-CTLA-4 and anti-PD-1 immunotherapeutic cohorts were not separately analyzed in each cancer type, which might have obscured the effects of sex (14, 42).

PD-1 blockade is effective to only a minority of melanoma patients, and the mechanisms are unclear (8, 9, 43). Note that in the previous studies, the gender factor has been ignored (8, 9, 22, 23, 26, 43). Importantly, for melanoma patients treated with anti-PD-1 therapy, we showed that high TMB correlated with durable clinical benefits only in males, providing a partial explanation for the heterogeneity in the patient responses.

One limitation of our study lies in the small number of PD-1 blockade samples from different clinical trials, but our datasets are much more homogeneous than previous studies (8, 9, 22, 23, 26, 43) and thus more reliable for discovering the determinants of therapeutic efficacy. In addition, we recruited multidimensional TCGA-SKCM datasets to corroborate our analysis. Although a recent study based on a melanoma immunotherapeutic cohort (22) also revealed sex-based TMB for predicting survival (HR: hazard ratio; females: HR=0.6, *P*=0.1; males: HR=0.49, *P*=0.005), we abandoned this cohort because it was based on targeted panel sequencing (MSK-IMPACT) and multiple ICB therapies.

Taken together, our study was the first to explore sex-related differences of TME in SKCM, revealing a significant positive link of males with high TMB to IFN-γ signaling, MHC-I gene expression, CTLs infiltration, tumor immunogenicity, and better anti-PD-1 response in SKCM.

## Abbreviations

SKCM: Skin cutaneous melanoma
TMB: tumor mutation burden
TCGA: The Cancer Genome Atlas
TME: tumor microenvironment
MHC: major histocompatibility complex
CTLs: cytotoxic T lymphocytes
ICB: immune checkpoint blockade
WES: whole-exome sequencing
CR/PR: complete or partial response to anti-PD-1 therapy
SD/PD: stable disease or progressive disease to anti-PD-1 therapy

---

**Table S1.** Summary of clinical and genomic characteristics in Cell-SKCM and TCGA-SKCM cohorts.

|  | No. (%) |  |
| --- | --- | --- |
| Clinical cohorts | Cell-SKCM | TCGA-SKCM |
| No. of patients | 78 | 472 |
| Immunotherapy | Yes | No |
| Median age, years(range) | 58(27-84) | 58(15-90) |
| Gender |  |  |
| Male | 48(62) | 292(62) |
| Female | 30(38) | 180(38) |
| Histology |  |  |
| Cutaneous Melanoma | 78(100) | 472(100) |
| Treatment | anti-PD-1 | No |
| Best overall response(BOR) |  |  |
| Complete/partial response(CR/PR) | 31(40) |  |
| Stable/progressive disease(SD/PD) | 45(58) |  |
| Not evaluable | 2(2) |  |
| Mutation profiles | WES | WES |
| RNA-seq profiles | No | Yes |

**Table S2.** Table for univariate and multivariate logistic regression of TMB, neoantigen burden, and age on BOR in Cell-SKCM cohort.

BOR (as a binary variable) **Female**
| Model | Covariates | Odds Ratio | P value |
| --- | --- | --- | --- |
| Univariate | TMB | 1.21 | 0.8 |
|  | Neoantigen burden | 1.03 | 1 |
|  | Age | 1.01 | 0.8 |
| Multivariate | TMB | 1.23 | 0.8 |
|  | Age | 1.01 | 0.8 |

BOR (as a binary variable) **Male**
| Model | Covariates | Odds Ratio | P value |
| --- | --- | --- | --- |
| Univariate | TMB | 7.96 | **0.009\*\*** |
|  | Neoantigen burden | 8.01 | **0.01\*** |
|  | Age | 1.02 | 0.4 |
| Multivariate | TMB | 7.77 | **0.01\*** |
|  | Age | 1.01 | 0.85 |

**Figure S1.**
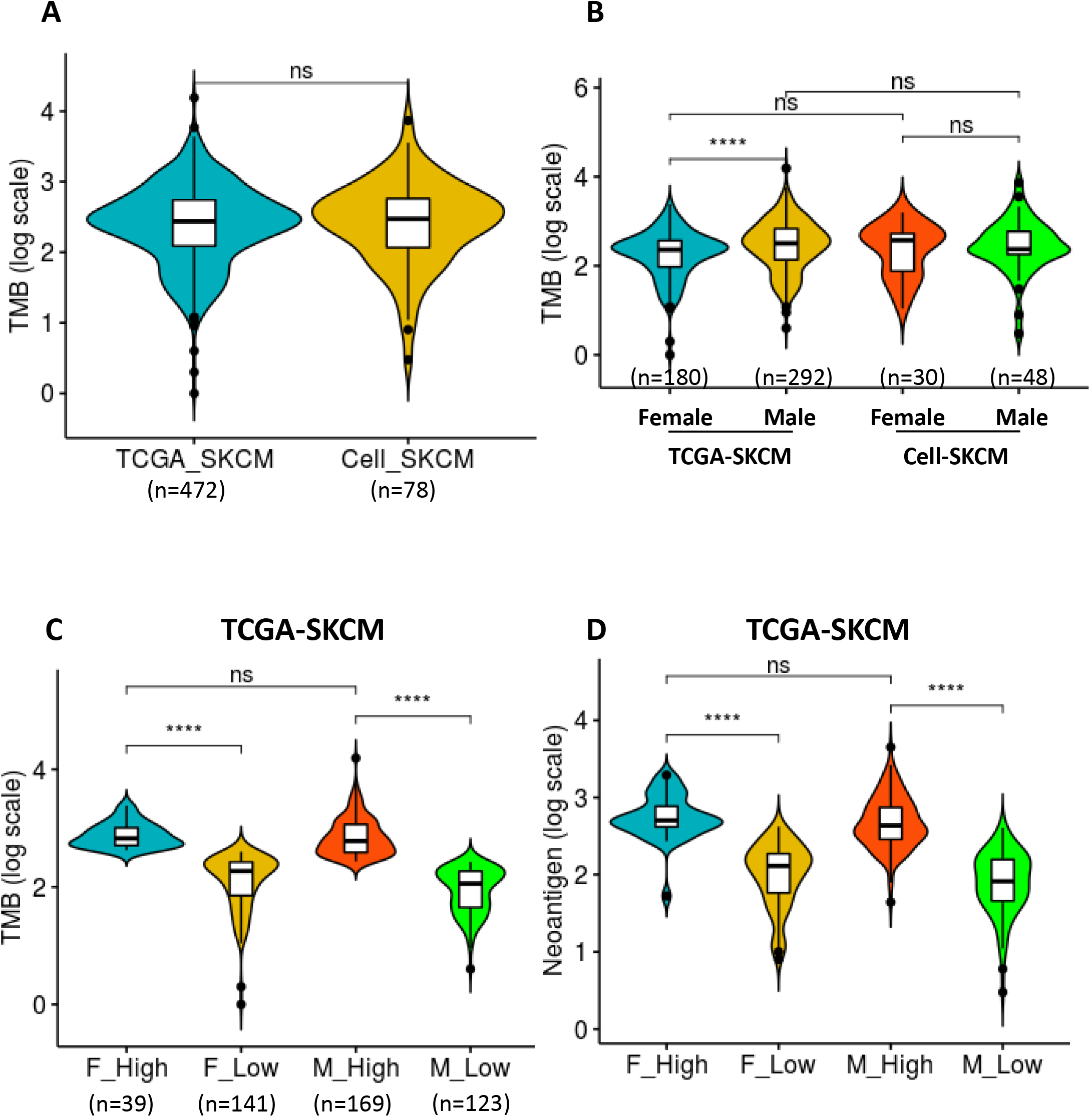

**Figure S2.**
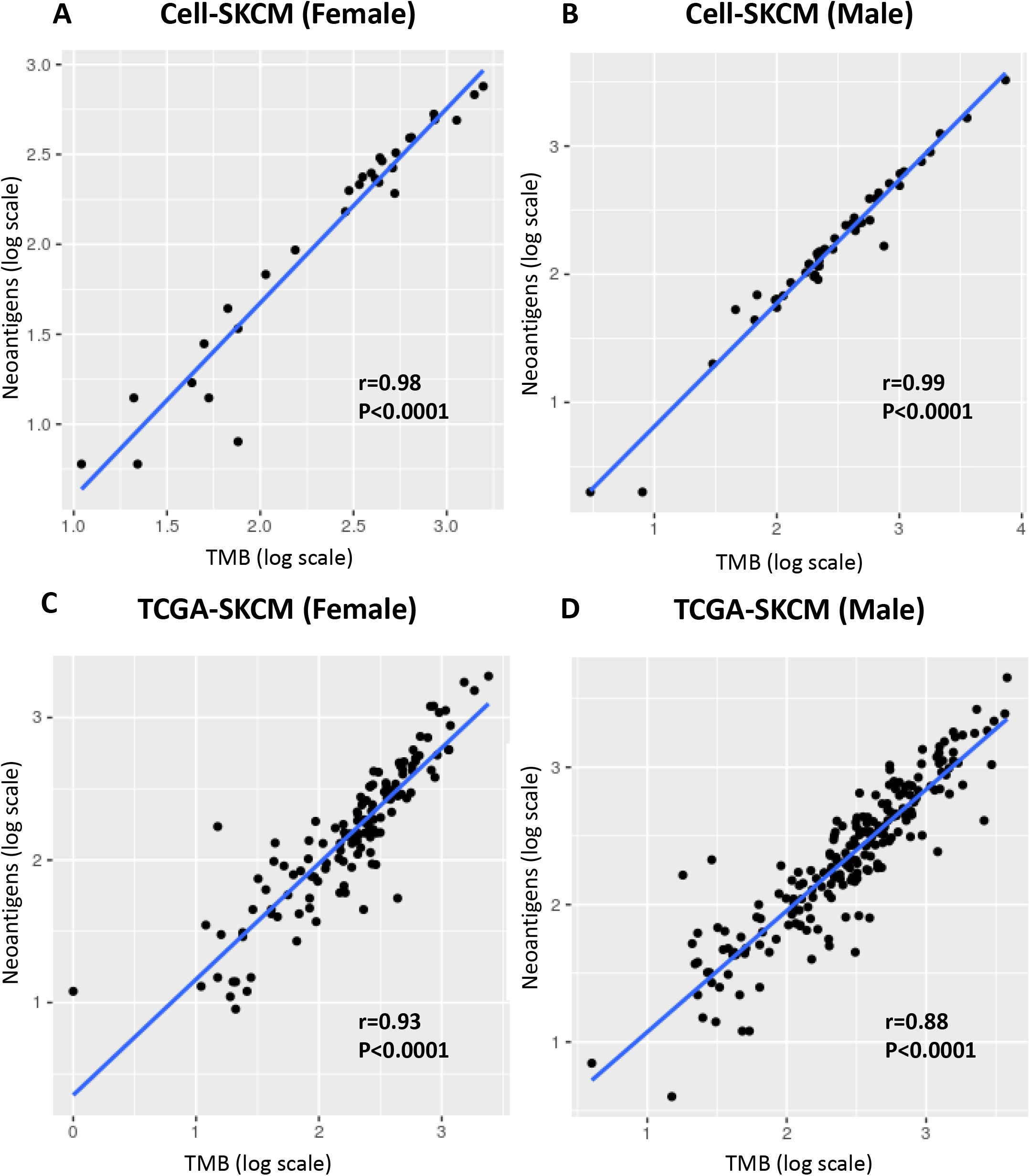

**Figure S3.**
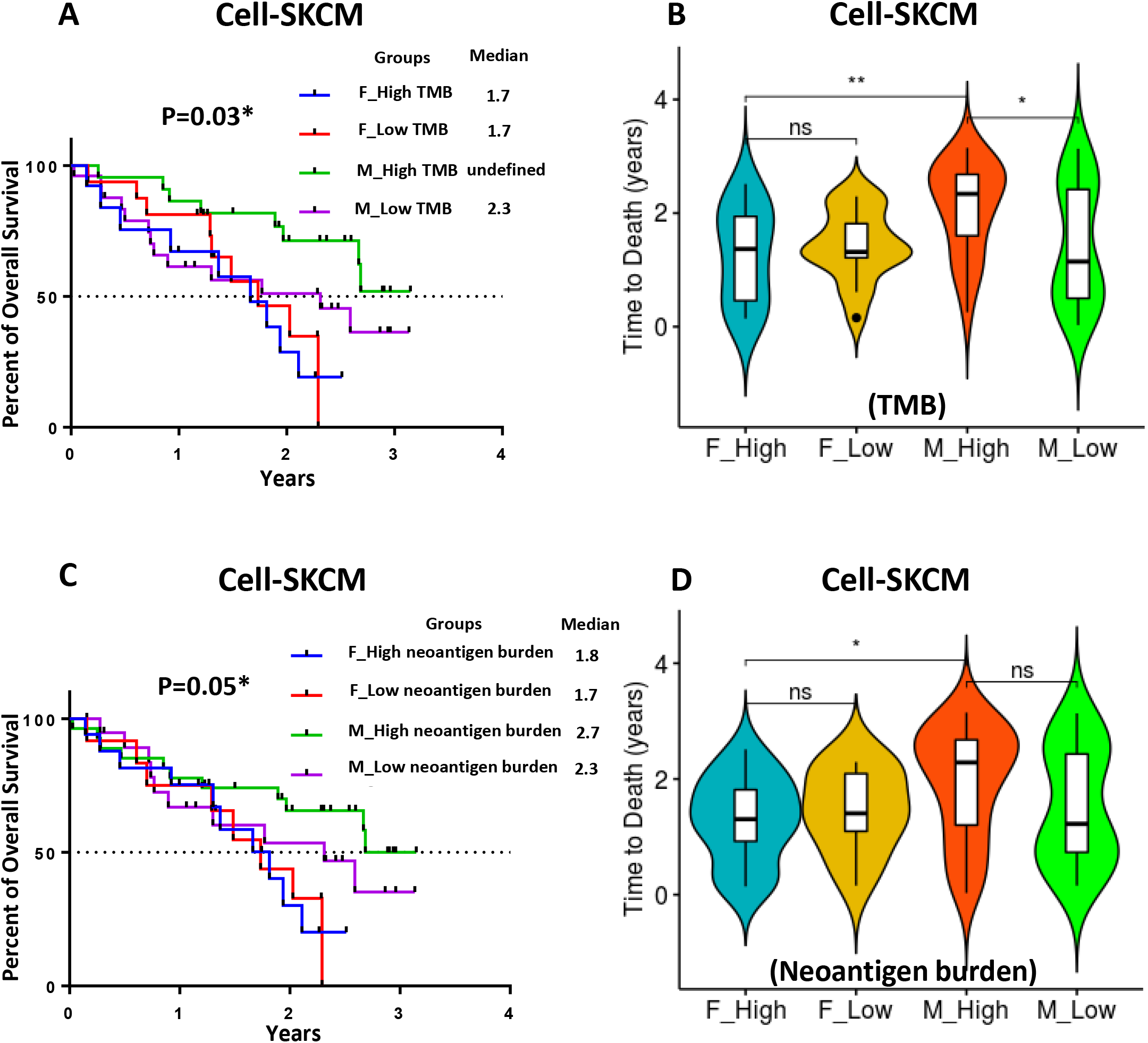

**Figure S4.**
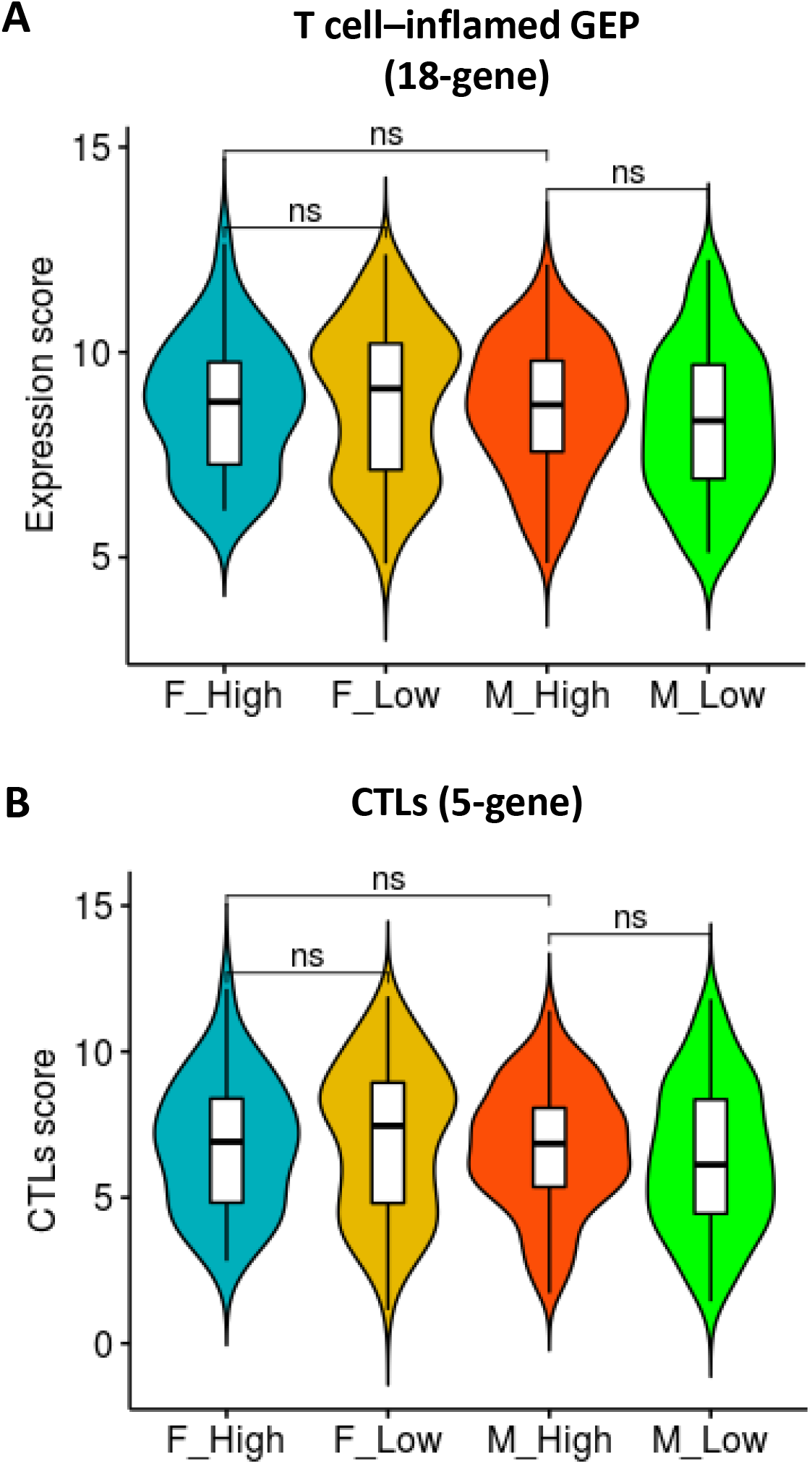

**Figure S5.**
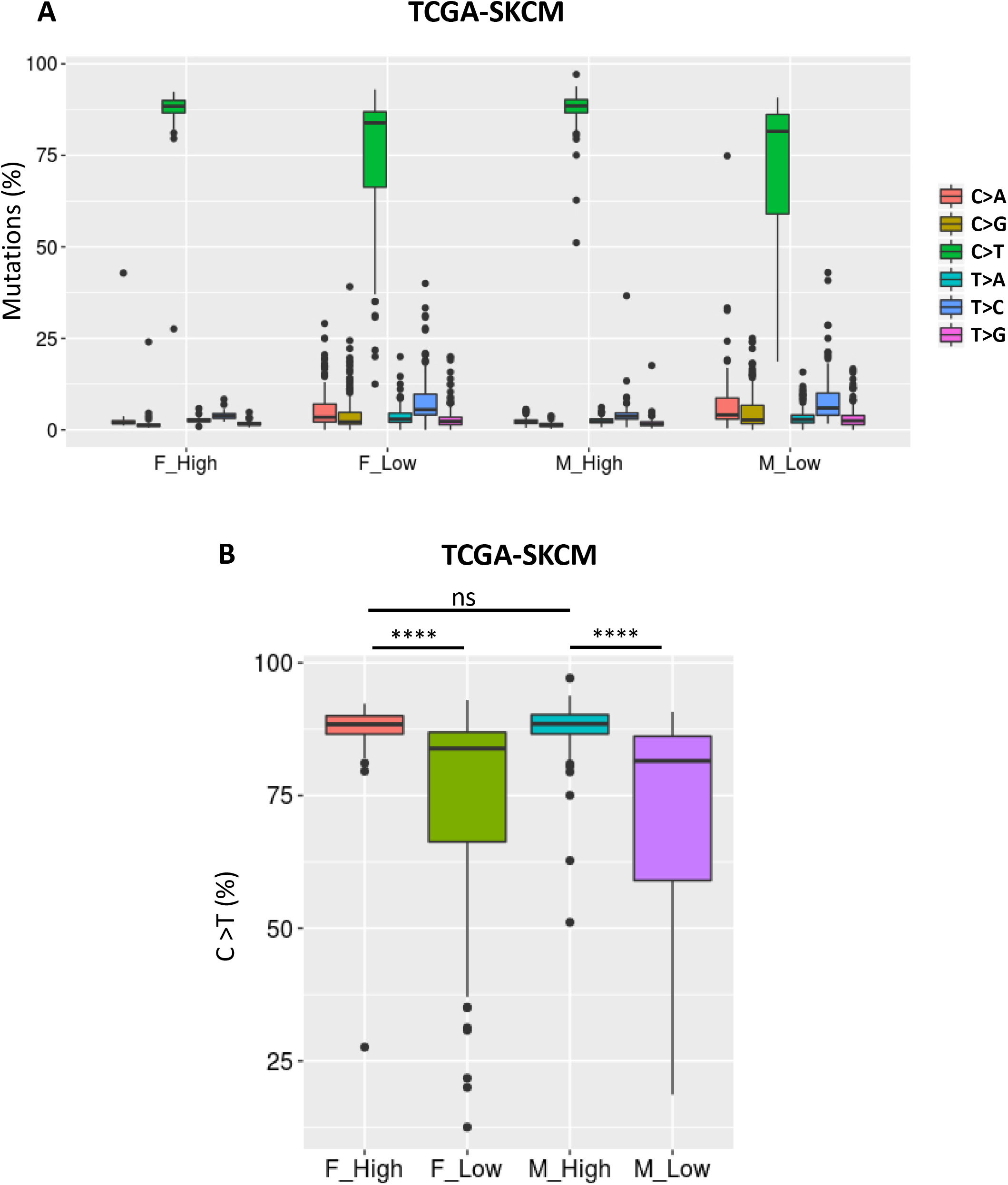

**Figure S6.**
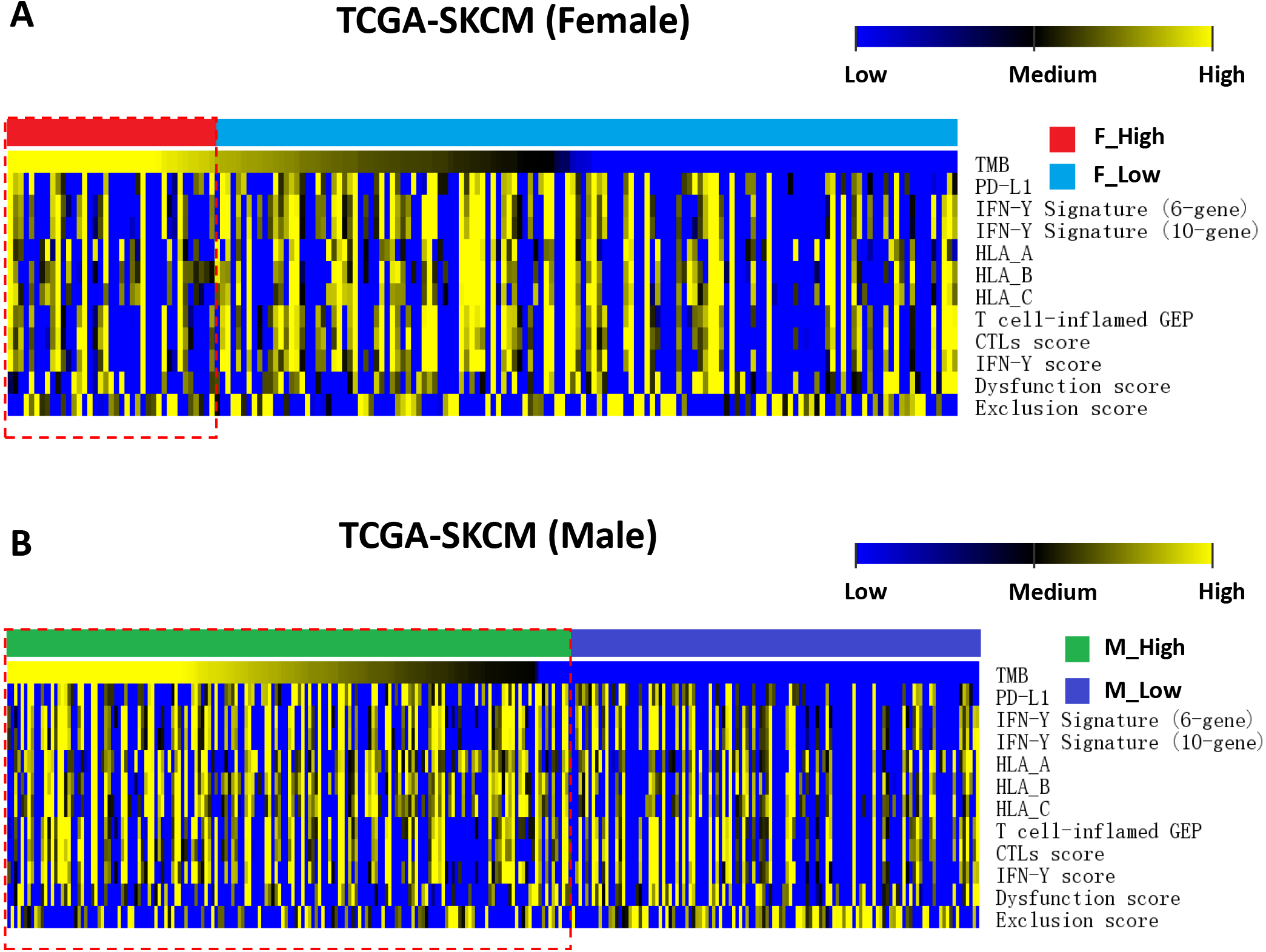

**Figure S7.**
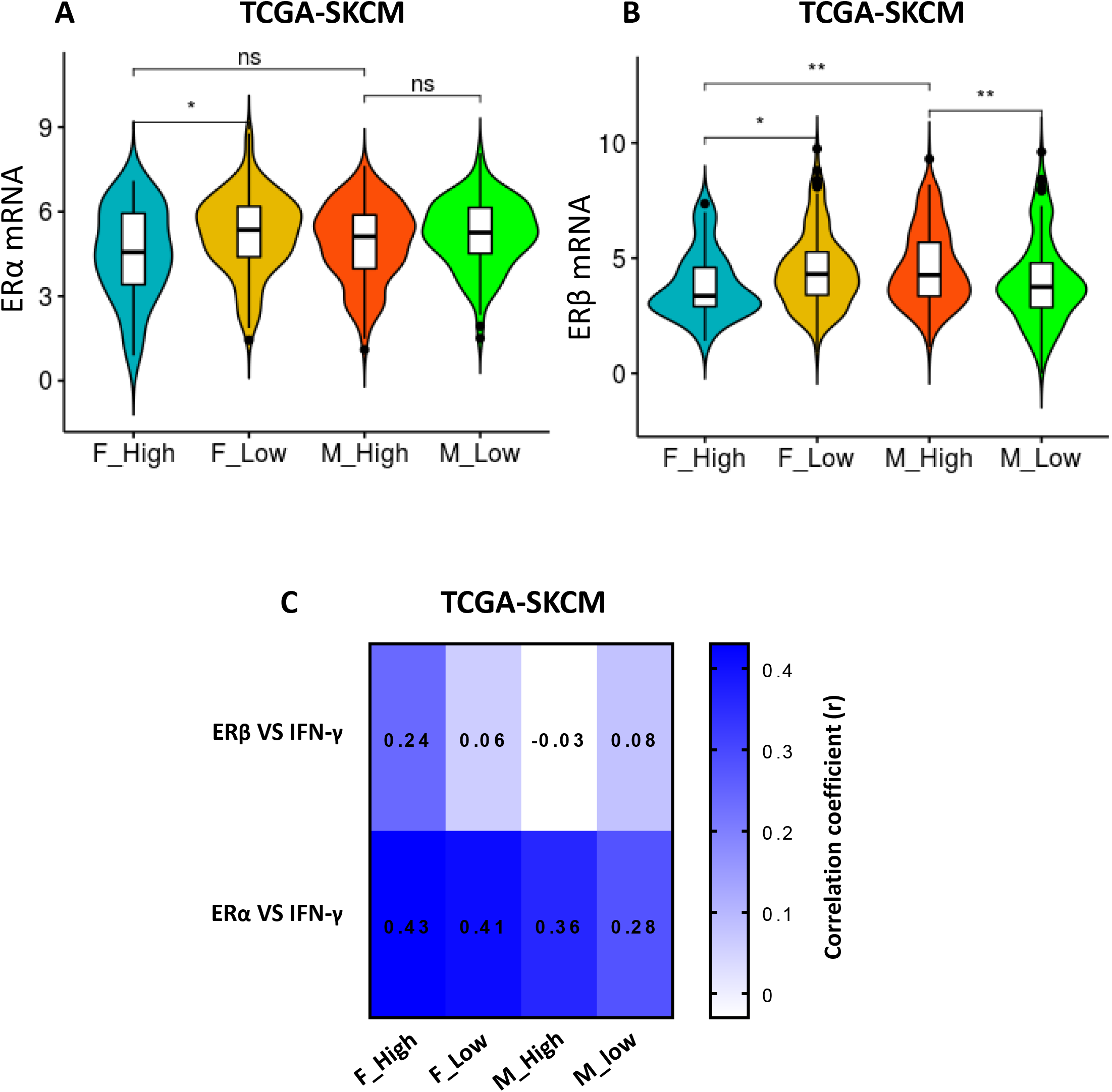

**Figure S8.**
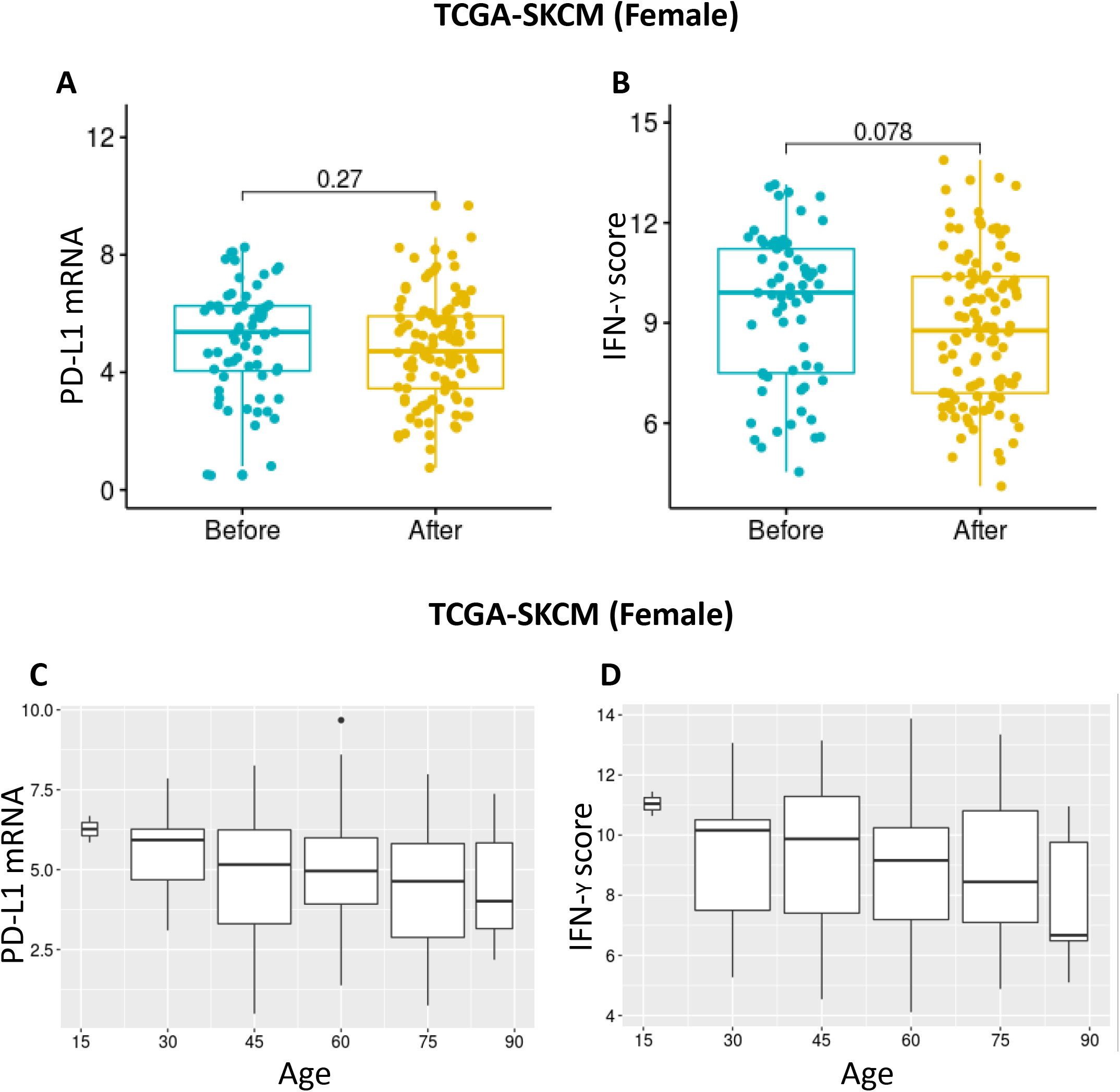

